# Cocaine Acts Through Sigma-1 to Enhance HIV Infection of Microglia

**DOI:** 10.1101/2025.08.06.668840

**Authors:** Oluwatofunmi Oteju, Xiaoke Xu, Taylor Kist, Katy Emanuel, Lexi Sheldon, Marzieh Daniali, Meng Niu, Howard Fox, Peter J. Gaskill

**Author notes:** Correspondence to: Peter Gaskill, Full address: 245 N 15^th^ Street, Philadelphia, PA, 19102.

## Abstract

Cocaine use disorder is highly comorbid in people with HIV and can accelerate infection, alter neuropathology, and exacerbate cognitive decline despite antiretroviral therapy (ART). Many of these effects are due to infection and dysregulation of CNS myeloid cells, especially microglia, which comprise a significant reservoir in this compartment. However, the precise mechanism(s) by which cocaine (Coc) enhances HIV infection in microglia are unclear, partly due to the lack of translationally relevant human microglial models suitable for mechanistic evaluation of Coc-mediated changes in viral dynamics. Canonically, Coc acts by blocking dopamine transporter activity, however, Coc has additional mechanisms beyond dopaminergic tone, involving the endoplasmic reticulum (ER) protein sigma-1, which has diverse cellular functions, including modulation of stress pathways such as the unfolded protein response (UPR). Viruses, including HIV, can exploit the UPR to amplify stress-induced protein production in host cells, enhancing viral replication. Therefore, we hypothesized that Coc-mediated activation of sigma-1 increases HIV infection of microglia via activation of the UPR. Using human-induced pluripotent stem cell microglia (iPSC-Mg), we evaluated changes in the percentage of infected iMg as well as p24Gag secretion using high-content imaging and AlphaLISA. Coc increased both p24 secretion and the number of infected iMg. The enhanced p24 secretion persisted despite ART, without the corresponding increase in percent p24, suggesting Coc augments the post-entry steps of viral replication. Inhibition of dopamine receptors did not diminish the impact of Coc, but pharmacological inhibition and CRISPR KO of sigma-1 blocked the effect. Independently, sigma-1 agonists increased p24 secretion, suggesting that Coc acts through sigma-1 rather than dopaminergic pathways. Single-cell RNA sequencing revealed distinct transcriptomic alterations in HIV+Coc-treated iPSC-Mg. Further genetic and proteomic validation confirmed activation of the IRE1-XBP1 branch of the UPR in the HIV+Coc condition, with increased XBP1 signaling and downstream cytokine (IL-4 and IL-7) secretion. The HIV+Coc condition also showed reduced expression of antiviral response genes and enhanced HIV transcriptional regulation genes. Immunofluorescence staining showed increased sigma-1 in p24-cells and reduced sigma-1 in p24+ cells in HIV+Coc cultures and revealed that HIV+Coc promotes sigma-1 movement to the ER. These findings suggest that Coc exploits sigma-1 signaling to modulate the UPR, enhancing viral replication and immune evasion. Sigma-1 emerges as a critical link between HIV-induced cellular stress and cocaine exposure, highlighting a shared molecular pathway that can be leveraged for the treatment of comorbid HIV neuropathogenesis, substance use disorder, and related neuropsychiatric disorders.

## Introduction

Substance misuse continues to increase globally, afflicting almost 300 million people in 2024^1^. It is particularly problematic in vulnerable populations, such as people with HIV (PWH), where it is highly comorbid, affecting 9 - 48% of PWH globally compared with 2% of the general population^1–3^. Among addictive drugs, cocaine (Coc), which is used by 5 - 15% of PWH, can disrupt initiation of and adherence to antiretroviral therapy (ART), as well as promote risky behaviors like needle sharing and unsafe sex, significantly increasing the spread of HIV^4–10^. Cocaine can also directly impact viral pathogenesis, significantly accelerating HIV progression in both the periphery^7,9,11–15^ and the central nervous system (CNS)^16–19^. The specific processes by which Coc influences HIV infection, particularly in the CNS, remain undefined, limiting our capacity to effectively treat HIV neuropathogenesis in PWH who use Coc.

HIV enters the CNS rapidly after initial infection, and can result in a constellation of neurological symptoms known as neuroHIV^20^. These issues remain prevalent in PWH despite effective ART^11,21,22^, suggesting that active HIV replication is not necessary to sustain neurological disease. The primary targets for HIV in the CNS are microglia and other CNS macrophages^22–25^. Astrocyte infection remains controversial, while neurons, oligodendrocytes, endothelial cells, and other CNS cells are not infected^26,27^. Microglia and other macrophages respond to HIV infection of the CNS by secreting inflammatory factors. HIV infection of these cells dysregulates them, further promoting a neuroinflammatory environment^28–30^. CNS myeloid populations may also house a viral reservoir in the CNS^24,25,31^, although the precise composition and viral replication status within this reservoir remain poorly understood^32^.

As Coc alone can also dysregulate microglial immune function and drive neuroinflammation^33–35^, the interaction of Coc and HIV could exacerbate neuropathogenesis. Coc has been shown to increase HIV infection in non-CNS macrophages^36–39^, although the effects of Coc on HIV infection in human microglia remain unclear. These data suggest that Coc use could amplify the number of dysregulated myeloid cells and/or expand the size of the CNS reservoir. Therefore, understanding the mechanisms driving Coc-mediated changes in HIV infection of microglia could inform targeted interventions for neuroHIV in PWH with cocaine use disorder (CUD).

Coc is canonically considered a dopamine transporter (DAT) inhibitor, preventing dopamine reuptake into the cell, but recent data show that Coc has other mechanisms of action beyond the modulation of dopamine. Notably, Coc can interact with sigma-1, a multifaceted protein embedded in the endoplasmic reticulum (ER) that is associated with a number of Coc-induced neurochemical changes^40,41^. Sigma-1 is expressed in multiple cell types, including macrophages and microglia^42–45^, and the majority of *in vitro* and *in vivo* studies indicate that it is neuroprotective^46,47^. However, in the context of HIV, sigma-1 can promote the formation of reactive oxygen species, inflammatory cytokine secretion, and increased microglial infection^42,48^. In the ER, elevated sigma-1 promotes the adaptive response against ER stress via the unfolded protein response (UPR)^49^, increasing the activation of cell survival pathways. The mechanism(s) and/or downstream mediators by which sigma-1 drives Coc-induced changes in viral dynamics are not well understood, but viruses, including HIV, can exploit the UPR to amplify stress-induced protein production in the host cells and increase viral replication^50–55^.

Our data show that Coc enhances HIV infection of human iPSC-derived microglia-like cells (iMg), increasing both the percentage of HIV infected (p24+) iMg and the secretion of p24, a surrogate for viral replication. This mirrors findings in other myeloid cell models^38,42,56–61^, but it is the first time this has been shown in iMg. This increase persisted even in the presence of the first-line antiretroviral therapy (ART) combination of bictegravir, emtricitabine, and tenofovir alafenamide (trade name Biktarvy), and pharmacological inhibition of dopamine receptors did not diminish the impact of Coc. However, inhibition of the sigma-1 did block the effect, indicating that activation of this receptor was central to the process. This was validated using multiple sigma-1 agonists and antagonists. Single-cell RNA sequencing (scRNAseq), validated by qPCR and Western Blot, revealed Coc enhances the expression of genes and proteins associated with the transcription factor X-box binding protein 1 (XBP1)-driven unfolded protein response and HIV transcriptional regulation. Exposure to Coc likewise reduces expression of genes and proteins associated with host antiviral response. A multiplex cytokine panel supported this, showing that cocaine treatment reduced antiviral cytokines and increased pro-survival cytokines. High content, confocal microscopy, and analysis of HIV-infected iMg cultures showed that Coc increased sigma-1 expression in uninfected (p24-) iMg, but not infected (p24+) iMg, and promoted sigma-1 localization to the ER. These data suggest that Coc-mediated changes in HIV replication could be driven by bystander cells with low or no HIV infection, as opposed to highly infected cells. These findings delineate a mechanism through which Coc enhances HIV infection in microglia, potentially via exploiting the innate immune and unfolded protein responses in bystander cells, promoting viral replication and potentially increasing susceptibility to infection.

## Methods

### Differentiation and Culture of human induced pluripotent stem cell microglia (iMg)

Human, microglia-like cells (iMg) were differentiated from induced pluripotent stem cells obtained from the Children’s Hospital of Philadelphia (CHOP) for wild-type (WT) iPSC and Jackson Lab for sigma-1 knockout (SIGMAR1 KO) iPSC as common myeloid progenitors (CMPs). This process uses an established 11-day protocol^55,62,63^ generating iMg that express common microglia identity markers, including P2RY12, TMEM119, and IBA-1. Briefly, CMPs are differentiated in RPMI-1640 supplemented with 10% FBS, 0.1% PenStrep, IL-34 (100ng/mL, PeproTech, 200-34-10UG), TGF-β (50ng/ml, PeproTech, 100-21-10UG), and M-CSF (25ng/ml, PeproTech, 300-25-10UG) at 37°C in a humidified incubator under 5% CO2. Half media changes were performed on days *in vitro* (DIV) 2 and 5, while full media changes were performed on DIV 8 and 11, and the cytokines IL-34, TGFβ, and M-CSF were added fresh with each media or ½ media change. At DIV11, cells were considered mature and ready for downstream assays. iMg were cultured in glass-like polymer plates (Cellvis, 96-well, P96-1.5P or 24-well, P24-1.5P, depending on the experiments) at a density of about 312.5 cells/mm^2^, 10,000 cells / well in 96-well plates, and 66,600-75,000 cells / well in 24-well plates. In this study, we included iMg derived from four distinct 4 iPSC lines (WT6, WT10, WT15, WT17).

### HIV Inoculation of iMg

Mature iMg plated in 96-wells were inoculated with HIV_ADA_ (1ng/ml, otherwise stated) for 24 hours, then washed to remove excess virus. For cocaine and agonist treatments studies, iMg were inoculated with HIV_ADA_ in the presence of cocaine (10^-5^ – 10^-8^M) or sigma-1 agonists (PRE-084 or Donepezil 10^-6^ – 10^-8^M). For antagonist studies, iMg were pretreated with antagonists against different targets (BD1063 10^-6^M: sigma-1 antagonist, Flupentixol 10^-6^M: pan-dopamine receptor antagonist) for 45 mins prior to inoculation with HIV_ADA_, media was replaced with fresh media containing virus. Following the wash at 24 hours, cultures were given fresh media, and supernatant was collected and replaced with fresh media every 48 hours up to 9 days post-infection (DPI). Supernatants from all wells of the same inoculation condition were pooled, aliquoted, and stored at −80°C for analysis of p24Gag as a measure of viral replication. For studies that included combination antiretroviral therapy (ART), the antiretroviral drugs Bictegravir (BIC) + Emtricitabine (FTC) + Tenofovir Alafenamide (TAF), which comprise the first-line regimen Biktarvy. These drugs were added to each well along with the fresh media at 3 DPI. The drugs were used at concentrations mimicking the peak plasma concentrations (C_max_) found in patients: BIC, 13.5 μM; TAF, 600 nM; FTC, 7 μM. Supernatants were analyzed for p24Gag using AlphaLISAs performed according to the manufacturer’s protocol (Revvity).

### Single Cell RNA Sequencing

#### Sample preparation

Mature iMg cultured in 24-well plates (Cellvis, P24-1.5P) were inoculated with HIV_ADA_ (20 ng/ml) for 24 hours, after which virus was washed off cells, and cells remained in culture till 5 days post-infection. On day 5 dpi, cells were treated with cocaine (10^-8^ M) for 3 hours. Following treatment, the cells were washed with PBS, then detached at 37°C using TrypLE (12604013, Thermo Fisher). Detachment was halted by the addition of an equal volume of warm media, and cells were spun at 400xg for 5 minutes. iMg was cryopreserved in 90% FBS 10% DMSO at a minimum concentration of 1,000,000 cells/mL at −80°C > 1 day (Mr. Frosty; Thermo Fischer Scientific), then transferred to a liquid nitrogen tank.

#### Library preparation

Post-sorting, isolates were concentrated to approximately 1000 cells per µL and counted on the LUNA-FL Dual Fluorescence Cell Counter using AO/PI (acridine orange/propidium iodide) for viability and concentration. The cells were processed using the 10× Genomics (Pleasanton, CA, USA) Chromium 3’ v3 GEM kit targeting 8000 cells. Based on the targeting cell number, the ideal volume of cells was loaded onto the 10× Genomics Chromium GEM Chip and placed into the Chromium Controller for cell capturing and library preparation. In short, this occurs through microfluidics and combining with Single Cell 3’ Gel Beads containing unique barcoded primers with a unique molecular identifier (UMI), followed by lysis of cells and barcoded reverse transcription of RNA, amplification of barcoded cDNA, fragmentation of cDNA to 200 bp, 5’ adapter attachment, and sample indexing as the manufacturer instructed with version 3 reagent kits. The final average library sizes are around 450- 470bp and were sequenced using Illumina (San Diego, CA, USA) Nextseq550 and Novaseq6000 sequencers. The sequences will be deposited in NCBI GEO (accession number: in progress).

#### Bioinformatic analysis

Sequenced samples were processed using the 10× Genomics Cell Ranger pipelines (v7.1.0). scRNA-seq data were demultiplexed and aligned to a customized GRCh38 human reference genome (NCBI RefSeq assembly), which included an additional chromosome representing the HIV genome. Details of the alignment, quality control, PCA analysis, and clustering processes are found in the supplementary material. To identify DEGs between vehicle (reference group) and cocaine-treated samples, a t-test was applied using sc.tl.rank_genes_groups. Genes with a statistical score of zero were excluded. Remaining genes were ranked by statistical score and used for gene set enrichment analysis (GSEA) using the gseapy package (v1.1.3)^64^. Enrichment was performed using the MSigDB Hallmark and Gene Ontology (Biological Process) gene sets, using minimum and maximum gene set sizes of 5 and 500, respectively. Significantly enriched gene sets were defined by an FDR q-value < 0.05.

### Statistical Analysis

Statistical analyses were performed using GraphPad Prism. Data distributions were assessed for skewness, normality, and lognormality using the Shapiro-Wilk, Kolmogorov-Smirnov, and D’Agostino & Pearson tests. Outliers, presumed to be technical artifacts, were identified and excluded using the ROUT test (Q = 0.1%). p24 AlphaLISA data were log10-transformed prior to analysis. Gene expression data were analyzed as 2^−ΔCT. Group comparisons were made using paired t-tests (for normally distributed data with equal variances) or Wilcoxon tests (for non-normal data). Multiple group comparisons were assessed using one-way ANOVA, two-way ANOVA, or mixed-effects analysis, followed by appropriate post hoc tests (Sidak’s, Dunnett’s, or Tukey’s) as necessary. Data are presented as mean ± SEM, with p < 0.05 considered significant.

## Results

### Characterization of HIV infection in iMg

As primary microglia are difficult to obtain, we used our established pluripotent stem cell-derived microglia-like cells (iMg) to evaluate HIV infection dynamics of microglia in the presence of Coc. Our prior work showed these cells exhibit microglia-like morphology and express the identity markers IBA-1, P2RY12, and TMEM119^62^. Mature iMg were inoculated with an R5-tropic strain of HIV (HIV_ADA_) at concentrations of 0.01ng•p24/mL – 20ng•p24/mL for 24 hours. Supernatants collected at 3, 5, 7, 9 days post-infection (DPI) were assessed for concentration of p24 as a surrogate for viral replication. Cultures were also fixed at the assay endpoint (DPI9) and stained for p24, nuclei, and cell bodies (**Fig.1A,B)** to assess the percentage of infected (p24+) iMg cells via high-content analysis. Analysis of p24 secretion demonstrated increasing secretion over time in response to all inoculating concentrations except the lowest (0.01ng•p24/mL), with the magnitude of secretion proportional to the concentration of inoculating virus (**Fig.1C**). Fixation and staining for p24 at DPI9 demonstrated p24+ cells in response to all inoculations except 0.01ng•p24/mL, with the percentage of infected cells also proportional to concentration of viral inoculum (**Fig.1D**). There were significant increases in p24+ iMg inoculated with 20ng•p24/mL HIV relative to other concentrations, and both infected iMg and p24 secretion were detected in response to inoculation with as little as 0.05ng•p24/mL HIV. These data are consistent with the level of HIV infection detected in the CNS of PWH using ART, with 0.5% of microglia containing cell-associated HIV RNA and HIV DNA^22^. Enumeration of iMg at DPI9 (**Fig.1D, blue line**) shows no change in the number of iMg in response to different inoculation, suggesting that changes in infection dynamics are not due to changes in cell viability. These data show that the iMg used in these studies are susceptible to infection and establish the assay window and limit of detection for these analyses.

**Figure 1:**
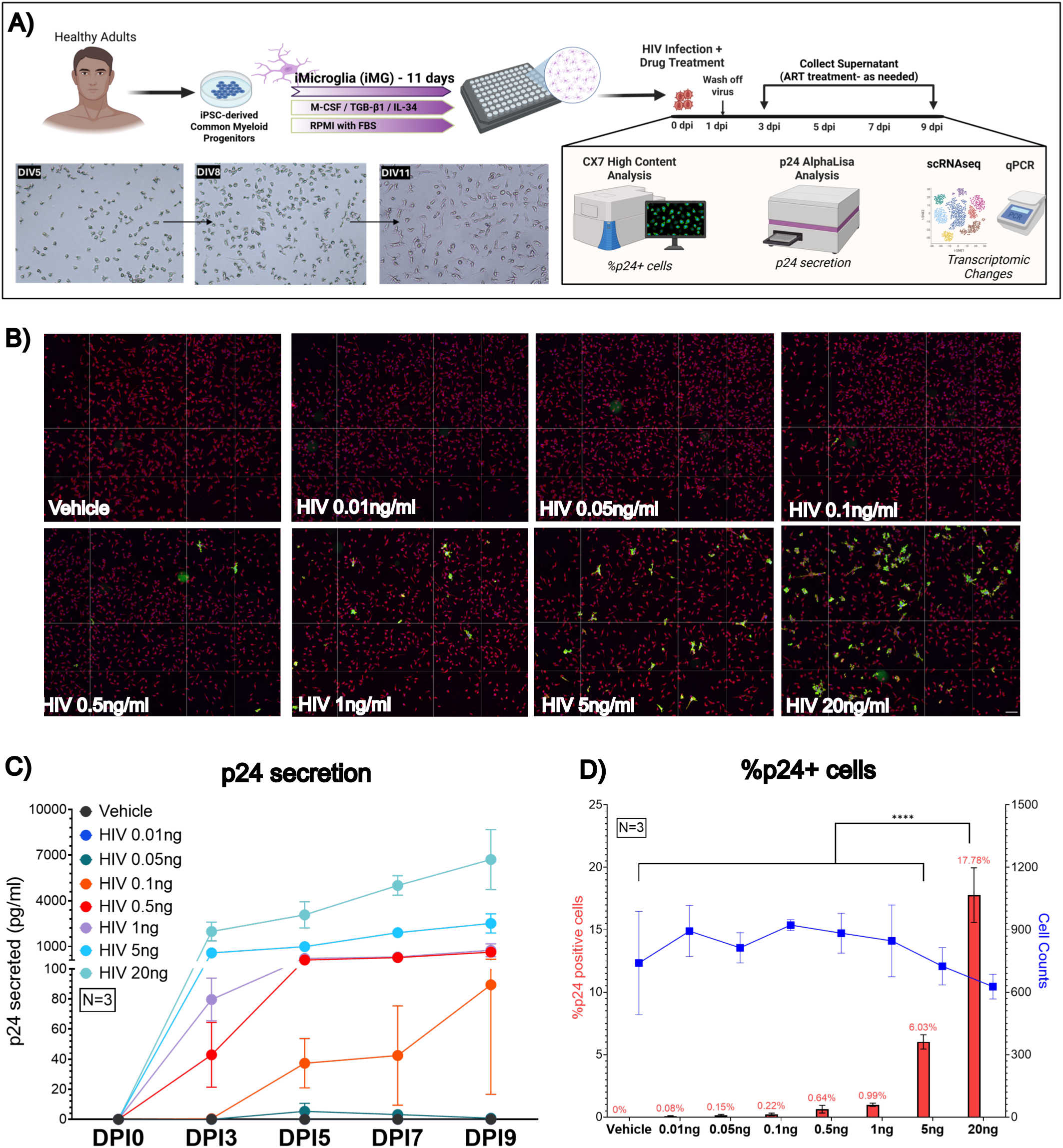
Validation of iMicroglia to study HIV viral dynamics. (**A**) iMg were differentiated from common myeloid progenotors (CMPs) treated with CSF1R-stimulating cytokines (IL-34 and M-CSF) and TGF-β1 for 11 days. HIV_ADA_ and drug treatments were performed for 24 hours, then supernatant collections were performed every two days till DPI9. **(B)** Mature iMg inoculated with HIV_ADA_ (0-20ng/ml) for 24hrs and imaged at DPI9. Cytoplasm (Red), Nucleus (Blue), p24Gag (Green), Scale bar = 20um. **(C)** p24AlphaLISA measuring virus secretion at DPI3, DPI5, DPI7, DPI9, showing dose-dependent increase in viral secretion. **(D)** Left axis (red bars): Percentage of p24+ cells at DPI9, Right axis (blue line): Cell counts (n=3, 3 iPSC donors). Two-way ANOVA [F (6, 12) = 57.6] followed by Dunnett’s test, HIV 20ng vs all concentrations ****P < 0.0001. Scale bar = 20um

### Cocaine increases HIV infection of iMg

To determine the optimal concentration of Coc for these studies, iMg were infected with HIV in the presence of a range of Coc concentrations (10^-8^ M-10^-5^ M, **Supplementary** Fig.1A) chosen to model the estimated CNS Coc levels needed to produce physiologic effects^65–68^. All Coc concentrations moderately increased p24 secretion over time relative to HIV alone, with greater increases at lower concentrations (10^-8^ M-10^-6^ M). Analysis of cell counts and LDH release (**Supplementary** Fig.1B**, C**) demonstrated that these differences were not driven by changes in iMg viability. The lowest concentration of Coc tested, 10^-8^ M, produced the greatest increase in p24 secretion, and *in vivo,* all microglia exposed to higher doses of Coc would also be exposed to this concentration due to degradation^69–71^, so 10^-8^ M Coc was used in subsequent experiments.

Human cells show substantial variability in susceptibility to HIV infection, so iMg from four distinct iPSC lines were infected with 1ng•p24/mL HIV. We assessed both p24 secretion over time and percentage of infected iMg at DPI9. Pooled results from multiple experiments across all four lines show that Coc (10^-8^ M) significantly increased both the percentage of infected (p24+) cells at DPI9 (**Fig.2A, B)** and p24 secretion at DPI3, 5, 7, 9 (**Fig.2C**). Non-linear regression analysis of p24 secretion shows that Coc significantly accelerates HIV infection of iMg (HIV *Slope* = 96.59, HIV+Cocaine *Slope* = 129.4; \**p* = 0.0453, *F* = 4,084, *df* = 1). To further examine donor variability, p24 secretion was separated by donor, showing that Coc induces some increase in p24 secretion in all donors. iMg from donors WT6 and WT10 were the most responsive, showing increased p24 secretion at all timepoints, as well as increased percentage of p24+ iMg at DPI9 (**Fig.2D, E**). Cells from WT17 showed a moderate increase in HIV secretion in response to Coc (**Fig.2F**), while cells from WT15 had a mixed response to cocaine that varied with timepoint (**Fig.2G**). These data corroborate human heterogeneity in both susceptibility to HIV infection and response to addictive substances^72,73^, and confirm prior studies showing Coc increases HIV replication^38,57,61,74^, although this is the first study – to our knowledge – that shows this effect in human iMg.

**Figure 2:**
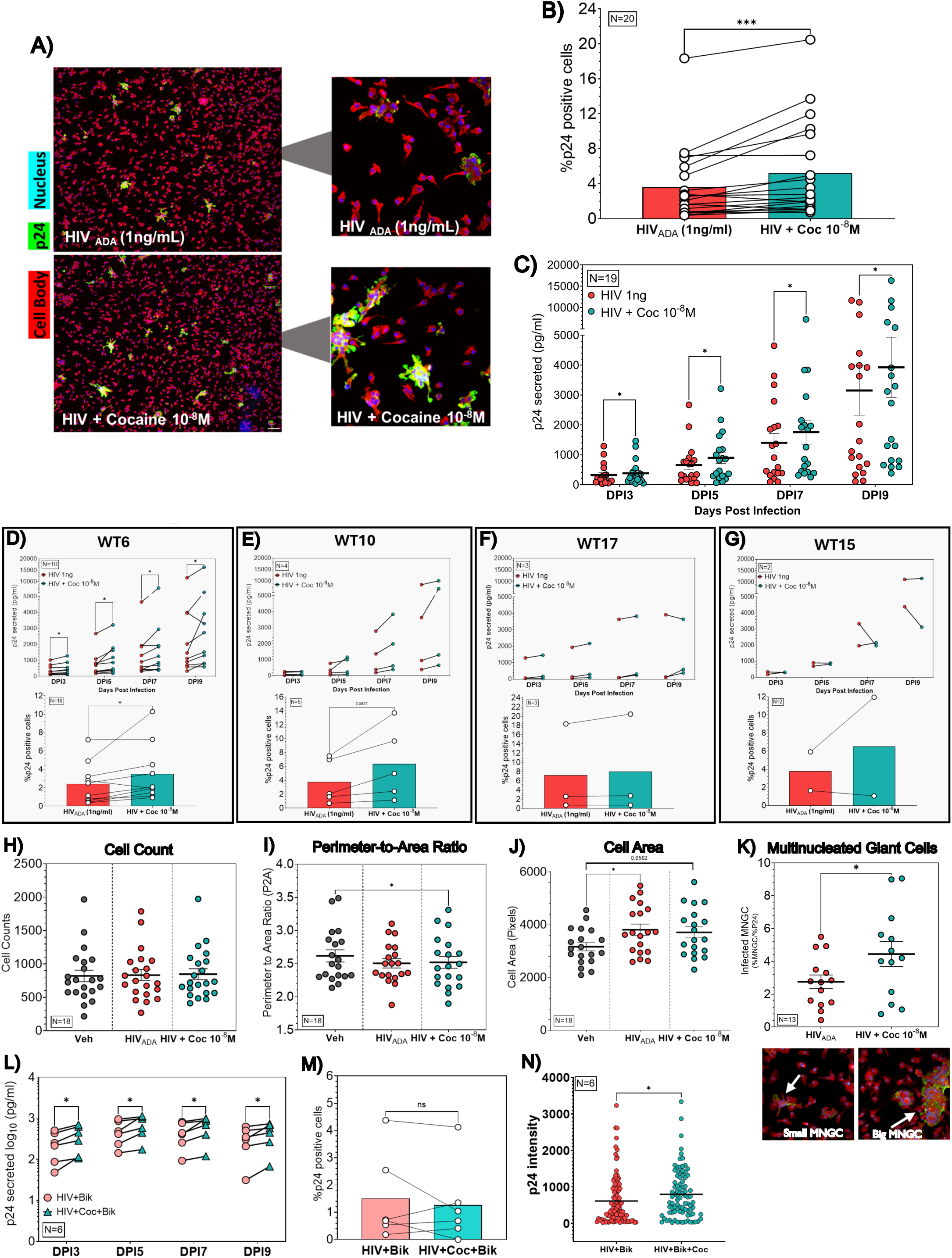
Cocaine increases HIV infection of iMg. **(A)** iMg treated with cocaine (10^-8^M) imaged at DPI9. Cytoplasm (Red), Nucleus (Blue), p24Gag (Green), Scale bar = 20um. **(B)** High content analysis quantification of %p24+ cells at DPI9 following cocaine treatment. N=20 (4 iPSC donors). Wilcoxon paired two-tailed test ***P < 0.001. **(C)** p24 AlphaLISA measuring cocaine-mediated increase in virus secretion at DPI3, DPI5, DPI7, and DPI9. Multiple Wilcoxon tests *P<0.01. **(D)** Donor-level data examining viral dynamics using p24 secretion and %p24+ cells. High content image analysis of cell-level data measuring **(H)** cell counts, **(I)** perimeter-to-area (P2A) ratio, **(J)** cell area, and **(K)** multinucleated giant (MNGC) cells. Multinucleated giant cells were quantified as cells with >5 nuclei. **(L)** iMg treated inoculated with HIV +/- cocaine were exposed to ART treatment every media change starting at DPI3. p24AlphaLISA was used to quantify virus secretion, and **(M)** high content screening was used for % p24+ cells. N=6 (3 iPSC donors), Multiple paired t-test *P < 0.05, Wilcoxon test ns: non-significant. **(N)** The p24 intensity of each cell was analyzed to validate the amount of virus production following cocaine treatment in ART-exposed cells. N=6 (3 iPSC donors). Mann-Whitney test *P <0.05. Scale bar = 20um

To further examine the effects of Coc on HIV-infected iMg, high-content screening was used to assess changes in cell parameters (i.e., cell counts and cell morphology) in response to Coc. Coc exposure did not alter cell counts relative to Vehicle or HIV cultures (**Fig.2H, Supplementary** Fig.1B). Coc-treated, HIV+ cultures did show a slight but significant decrease in perimeter-to-area ratio (**Fig.2I**), indicating that HIV + Coc induces a slightly rounder, as opposed to ramified morphology and suggesting enhanced microglia activation. HIV infection, with and without Coc, also increased iMg area, possibly due to the formation of multinucleated giant cells (MNGC) (**Fig.2J**). To assess this, we quantified the number of infected MNGC, defined as contiguous cell bodies containing > 5 nuclei. High content analysis found small and large MNGC in both HIV and HIV + Coc cultures, with a greater MNGC index (% MNGC / % p24+ cells) in HIV + Coc cultures relative to HIV alone (**Fig.2K**).

In the current era, the majority of PWH are on effective ART^75^, but Coc use can still increase both viral rebound and the risk of neurological deficits in these individuals^6–15^. This indicates that ART does not fully ameliorate the impact of Coc on HIV infection in the brain. To determine whether ART blocked the effect of Coc on HIV infection in microglia, iMg derived from 3 donors (WT6, WT10, WT15) were inoculated with HIV +/- Coc, then a subset of cultures were treated with ART after supernatant collection on DPI 3. This simulates the initiation of ART in people who have active HIV infection. The Coc-mediated increase in p24 secretion was sustained in the presence of ART (**Fig.2L**), but interestingly, there was no change in the percentage of infected iMg at DPI 9 (**Fig.2M**). Using high content imaging to measure the intensity of the p24 staining in individual iMg at DPI 9 from 3 infections, we found that Coc increases the p24 expression in infected iMg (**Fig.2N**). These data suggest that even when ART decreases the spread of infection by inhibiting infection of new iMg, Coc increases the translation of p24 in iMg that were infected prior to the addition of ART.

### Sigma-1 drives the cocaine-mediated increase in HIV infection

Classically, Coc acts by blocking the activity of the dopamine transporter (DAT), preventing dopamine reuptake into the cell^76,77^. Coc has other mechanisms of action beyond the modulation of dopamine. Notably, Coc interacts with sigma-1, a multifaceted protein embedded in the ER. This interaction has emerged as a focal point in understanding Coc-induced neurochemical changes^40,41^. To examine whether sigma-1 activity drives Coc-mediated changes in HIV infection, iMg from four iPSC lines were inoculated with HIV in the presence of either Coc or the sigma-1 agonist PRE-084. Both Coc and PRE-084 increased the percentage of DPI9 p24+ cells **(Fig 3A, B)** and increased p24 secretion at DPI3, 5, 7, 9 (**Fig. 3C**). This was validated in a separate set of infections with Coc or Donepezil, an FDA-approved sigma-1 agonist for dementia treatment. Donepezil treatment during inoculation also mirrored the effects of Coc on the viral dynamics (**Fig. 3D, E**). To confirm the involvement of sigma-1, iMg from all donors were pretreated with sigma-1 antagonist, BD-1063, for 45 min before HIV inoculation and Coc treatment. BD-1063 treatment blocked the Coc-mediated increase in the percentage of p24+ and p24 secretion but did not affect HIV infection in the absence of Coc (**Fig. 3F, G)**. We further confirmed this using an iMg which did not express sigma-1 after CRISPR-mediated gene deletion (SIGMAR1 CRISPR KO iMg), which verified that the cocaine-mediated increase in p24 secretion was lost in KO microglia **(Fig. 3H, Supplemental Fig. 1D)**. We then examined whether Coc could influence HIV replication via activation of microglial dopamine receptors. The dopaminergic impact of Coc is thought to be mediated primarily via interaction with dopaminergic neurons, and data suggesting microglial autocrine signaling via secreted dopamine are controversial^78–82^. However, we and others have shown that myeloid cells express the machinery needed to synthesize, release, take up, and respond to dopamine and that the myeloid dopaminergic system can regulate HIV infection^62,63,83–85^. Likewise, sigma-1 has been reported to form complexes with dopamine receptors, suggesting sigma-1 modulation of dopamine receptor activity^86,87^. Therefore, iMg were pretreated with pan-dopamine receptor antagonist Flupentixol (Flux) for 45 min prior to HIV inoculation and Coc treatment. Blocking dopamine receptor activation with Flux had no effect on the Coc-mediated increase in HIV infection (**Fig. 3I, J**), suggesting the effects of Coc on HIV replication are specific to sigma-1 activation and independent of the dopamine receptor activity.

**Figure 3:**
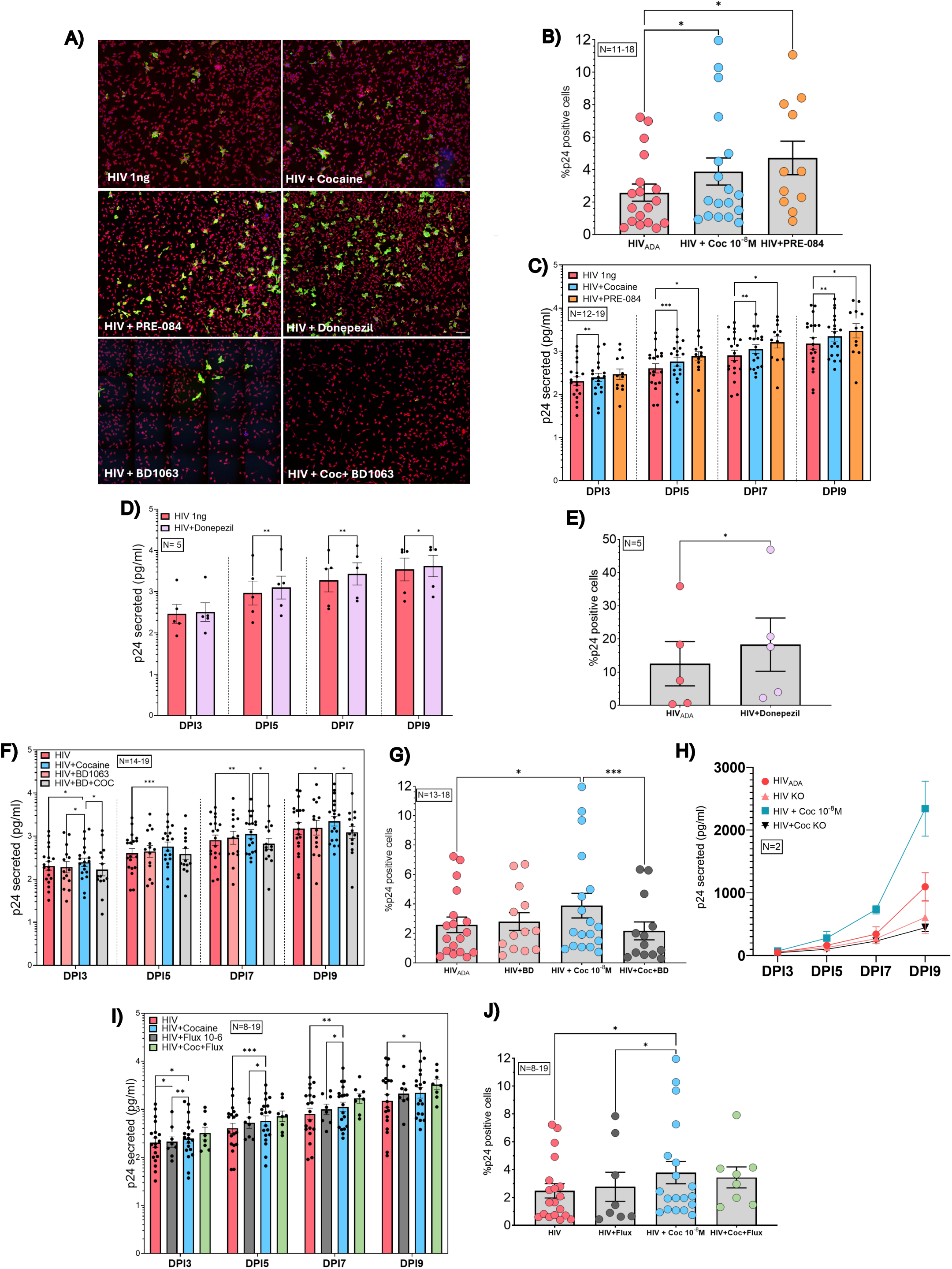
Sigma-1 drives the cocaine-mediated increase in HIV infection of iMg. **(A)** iMg were inoculated with HIV and cocaine (10^-8^M) or sigma-1 agonist (PRE-084 or Donepezil, 10^-6^M/10^-8^M, 24 hours), then evaluated for p24 secretion and % p24+ cells. **(B, C)** <u>PRE-084</u>: p24 secretion-Two-way Mixed effects analysis [F (0.8418, 15.15) = 25.14] followed by Dunnett’s post hoc test. %p24+ cells-One-way Mixed effect analysis [F (1.462, 19.74) = 4.965] followed by Dunnett’s post hoc test. **(D, E)** <u>Donepezil</u>: p24 secretion-Two-way ANOVA [F (3, 12) = 18.02] followed by Holm-Sidak post hoc test. %p24+ cells-Paired t test. **(F)** Pharmacological inhibition of sigma-1 (BD1063, 10^-6^M, 45min pretreatment before cocaine treatment) blocked cocaine mediated increase in p24 secretion and **(G)** % p24+ cells. p24 secretion-Two-way Mixed effects analysis [F (0.6640, 11.95) = 6.481] followed by Tukey’s post hoc test. %p24+ cells-Paired t test. **(H)** Sigma-1 CRISPR KO corroborated these studies. **(I)** iMg were treated with dopamine receptor antagonist (Flupentixol, 10^-6^M, 45min pretreatment before cocaine treatment), then evaluated for p24 secretion and **(J)** % p24+ cells. p24 secretion-Two-way Mixed effects analysis [F (0.6640, 11.95) = 6.481] followed by Tukey’s post hoc test. %p24+ cells-One-way Mixed effect analysis [F (3, 31) = 4.769] followed by Tukey’s post hoc test. *P<0.05, **<0 0.005, ***P<0.0005. Scale bar = 20um

### Cocaine modulates innate immune and XBP1-driven unfolded protein response

To explore targets and pathways altered in response to cocaine, single-cell RNA sequencing (scRNA-seq) was performed on DPI9 HIV-infected iMg treated with either vehicle (H_2_O) (HIV) or cocaine (HIV+Coc) for 3 hrs (**Fig.4A**). In total, 8 different cell clusters were identified (**Fig.4B**). Following quality control, doublet removal, and dimensionality reduction, there was a moderate segregation between HIV vs. HIV+Coc cells (**Fig. 4C**). HIV was one of the top upregulated DEGs with cocaine treatment, and all clusters showed increeased HIV expression with cocaine treatment, with differences in magnitude within clusters (**Fig. 4D, E, F**). For each cluster, we examined the percentage of cells altered between the HIV and HIV+Coc groups. The number of cells in clusters increased approximately 2-fold in the HIV+Coc samples, while clusters 0, 1 and 7 showed a 1.4-fold decrease, and clusters 4,5, and 6 had < 1.4-fold change with cocaine treatment (**Fig. 4G**). Given the high diversity of microglia, we mapped our identified clusters onto established microglia states^88–90^. Our analysis classified the clusters as follows: Disease-associated microglia (DAMs) and Neurodegenerative microglia predominantly in Cluster 2, with related populations in Clusters 4, 5, and 7; Homeostatic microglia in Cluster 1; Activated Response microglia in Clusters 0 and 2; and Proliferating Microglia in Cluster 6 **(Fig 4H)**. A total of 37 genes were upregulated in HIV+Coc relative to HIV, while 407 genes were downregulated. To better understand the impact of these cocaine mediated changes in HIV-infected iMg, Gene Ontology (GO, Biological Processes) and Molecular Signatures Databases (MSigDB) were used (Supplementary Table 5 and 6). Among the dysregulated pathways were UPR, oxidative phosphorylation, E2F targets, DNA repair, regulation of innate immune response, and interferon alpha response **(Fig. 4I, J)**, suggesting that cocaine modulates cellular stress responses to enhance HIV infection, possibly via sigma-1.

**Figure 4:**
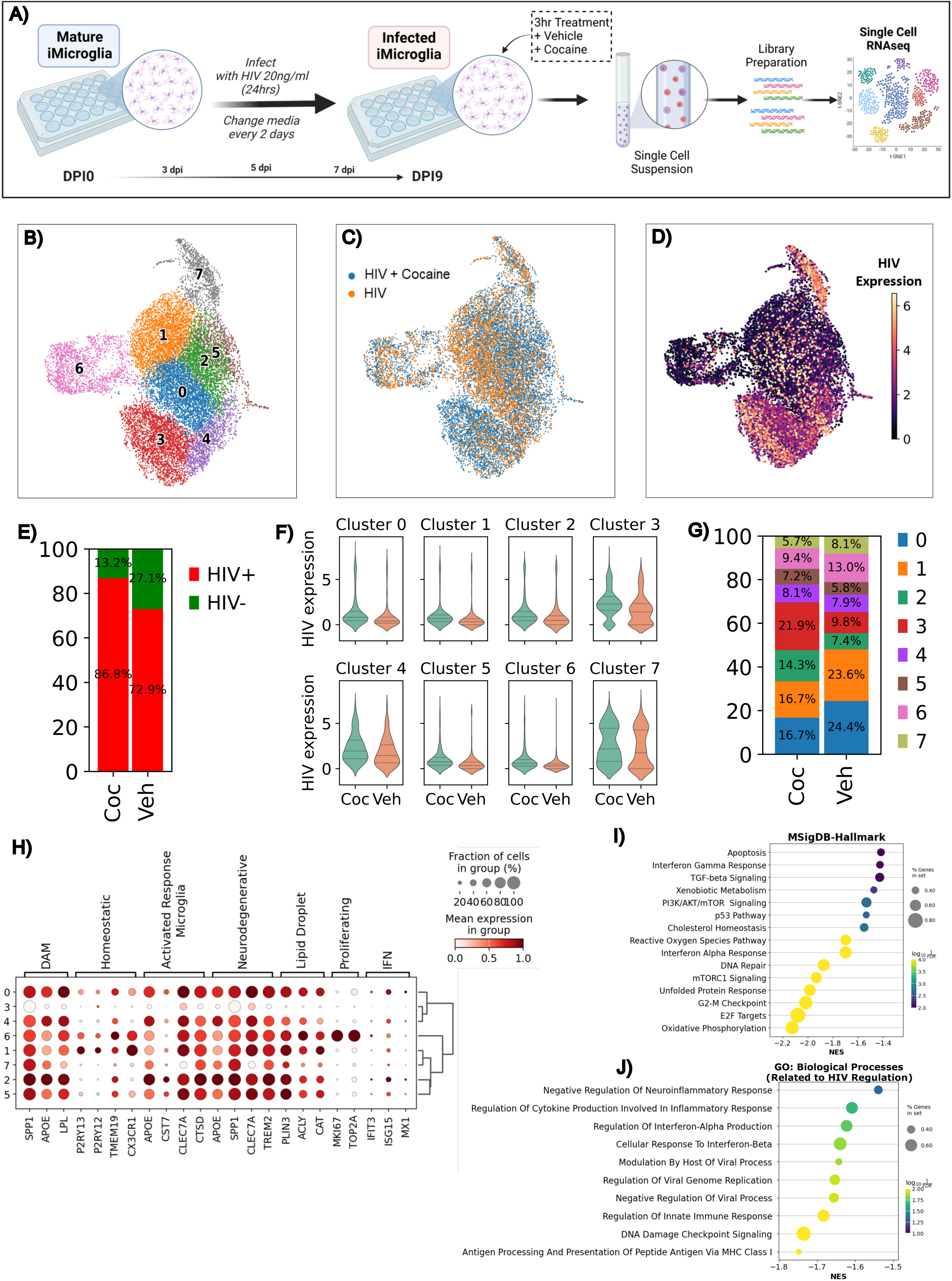
scRNAseq highlighting cocaine mediated transcriptomic changes in HIV infection of iMg. **(A)** Infected iMg treated +/- cocaine (10^-8^M) for 3 hours were lifted and prepared for scRNAseq analysis. **(B)** UMAP projection of 14,968 RNA expression profiles, colored by graph-based clustering results. **(C)** UMAP projection of single cells in the dataset, orange representing HIV treated cells and blue representing HIV + Cocaine treated cells. **(D)** UMAP projection of the range of HIV gene expression within single cells in the dataset. Darker colors indicate less HIV expression, while lighter colors indicate more HIV expression. **(E)** Bar graph showing changes in the proportion of HIV+ vs HIV-cells following cocaine treatment. **(F)** Violin plot showing cluster-level changes in HIV gene expression, with enhanced HIV expression in all clusters. **(G)** Bar chart showing the change in the proportion of cells in each cluster between HIV and HIV + Cocaine. **(H)** Markers associated with diverse microglia states within each cluster. Disease-associated microglia (DAM): *SPP1, APOE, LPL*; Homeostatic microglia: *P2RY12, P2RY13, TMEM19, CX3CR1*; Activated Response Microglia: *APOE, CST7, CLEC7A, CTSD*; Neurodegenerative: *APOE, SPP1, CLEC7A, TREM2*; Lipid-droplet associated microglia: *PLIN3, ACLY, CAT*; Proliferating: *MKI67, TOP2A*; Interferon-responsive microglia (IFN): *IFIT3, ISG15, MX1*. Gene set enrichment analysis using **(I)** Molecular signature database-Hallmark and **(J)** Gene ontology: Biological Processes.

Changes in each pathway were validated by qPCR and Western blot of select genes and proteins. For the UPR, cocaine enhanced XBP1 gene expression without changing genes that regulate other UPR pathways such as ATF6, ATF4, and CHOP (**Fig. 5A-D**). Sigma-1 modulates the UPR by stabilizing key signaling proteins such as IRE1α, upstream of XBP1, to maintain ER homeostasis, promote protein folding, and cell survival under stress. Western blotting of spliced XBP1, its active isoform, confirmed its increase at the protein level (**Fig. 5E, F**). Pathway analysis supported the validation of genes involved in regulation of the host innate immune response (↓IFIT1, IFIT2, IFITM1) and HIV transcriptional regulation (↑STMN1, ↓MALAT1) (**Fig. G**, **Supplementary** Fig. 2). Interestingly, the HIV+Coc group also showed a downregulation of interferon-induced transmembrane protein 1 (IFITM1), **Fig. 5H**, which restricts viral entry into cells^91,92^ and an upregulation in metastasis-associated lung adenocarcinoma transcript 1 (MALAT1, **Fig. 5I**), a long non-coding RNA that is associated with increased HIV transcription^93,94^. As the unfolded protein response regulates downstream cytokine release, cytokine secretion in HIV and HIV+Coc iMg was examined using a neuroinflammation multiplex cytokine panel. The secretion of IL-7 and IL-4 were increased in HIV+Coc cultures **(Fig. 5J, K**), while the secretion of IL-16, IL-12, and IL-6 decreased, confirming impaired antiviral response **(Fig. 5L, M, N)**, similar to patient data^37,95,96^. The UPR, specifically the XBP1 pathway, has been shown to regulate both IL-6 and IL-7^97,98^, and upregulation of IL-4 are is associated with XBP1 expression and activity^99^, further confirming the impact of HIV+Coc on the XBP1 UPR pathway.

**Figure 5:**
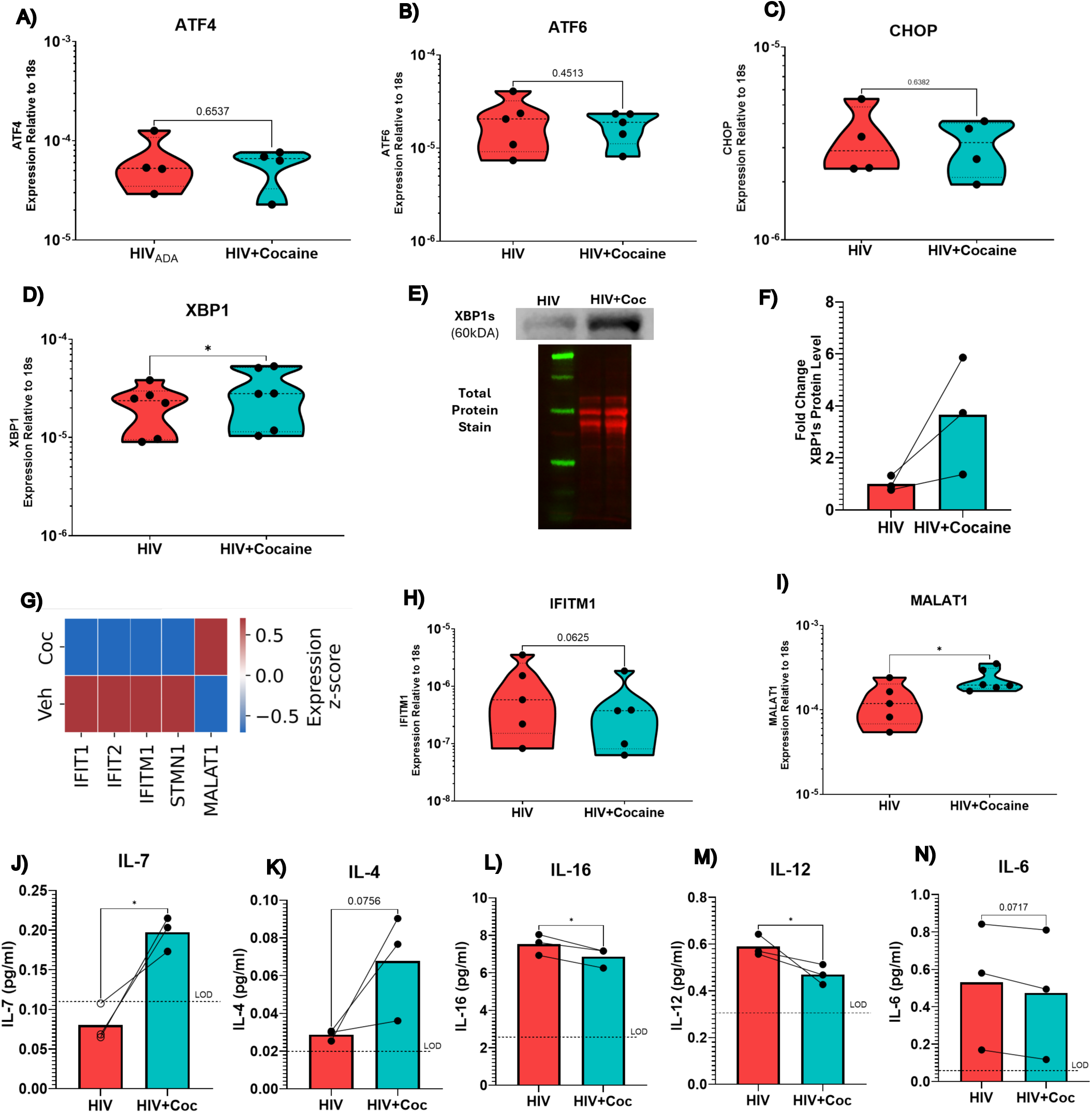
Cocaine modulates innate immune and XBP1-driven unfolded protein response. **(A-D)** qPCR gene expression (violin plot) validation of unfolded protein response genes: ATF4, ATF6, CHOP, XBP1. **(E, F)** Protein expression (bars) of XBP1s increased with HIV + Cocaine. **(G)** Single-cell RNA sequencing expression of antiviral (*IFIT1, IFIT2, IFITM1)* and HIV transcriptional regulation genes (*STMN1, MALAT1)*. qPCR gene expression validation of interferon gene **(H)** IFITM1 and **(I)** HIV transcriptional regulation gene, MALAT1. **(J-N)** Multiplex analysis of cytokines related to UPR and Innate immune response. Paired t-test. *P<0.05, **<0 0.005, ***P<0.0005

### Cocaine increases the expression of sigma-1 in HIV-infected cultures

When sigma-1 is activated in response to ER stress or exposure to an agonist, it translocates to different cellular compartments to mediate a variety of activities. Among these are the activation of the UPR in the endoplasmic reticulum (ER)^100,101^. To study the changes in sigma-1 expression and localization in response to HIV +/- Coc, uninfected and HIV-infected iMg were treated with either Veh (H_2_O) or Coc for 3 hr at 9 DPI, using a 3 hr treatment with the sigma-1 agonist PRE-084 as a positive control for sigma-1 binding. High-content immunofluorescence analysis showed sigma-1 intensity in the pattern HIV+Coc > HIV >> PRE > Coc > Veh (**Fig. 6A**). Each cell was then categorized as sigma-1^high^ or sigma-1^low^ based on immunofluorescence intensity. The designation of high or low was made by combining the intensity values across all treatments and defining anything above the 75^th^ percentile of the combined intensity as high, with everything else as low. A Chi-square analysis confirmed that the HIV+Coc condition had a significantly higher proportion of sigma-1^high^ iMg compared to HIV (**Fig. 6B**). To examine the changes in HIV and HIV+Coc iMg cultures in more detail, the HIV-infected (p24+) and uninfected (p24-) iMg within the same well were segregated into discrete populations and analyzed using high content screening. Interestingly, in both the HIV and HIV+Coc cultures, the uninfected (p24-) cells showed the highest level of sigma-1, with the uninfected (p24-) cells in the HIV+Coc culture showing significantly greater sigma-1 expression than either the HIV+Coc infected (p24+) cells or the HIV (p24-) cells **(Fig 6C)**. Additionally, in both the HIV and HIV+Coc cultures, the proportion of sigma-1^high^ iMg was greater in the uninfected (p24-) population (**Fig 6D**).

**Figure 6:**
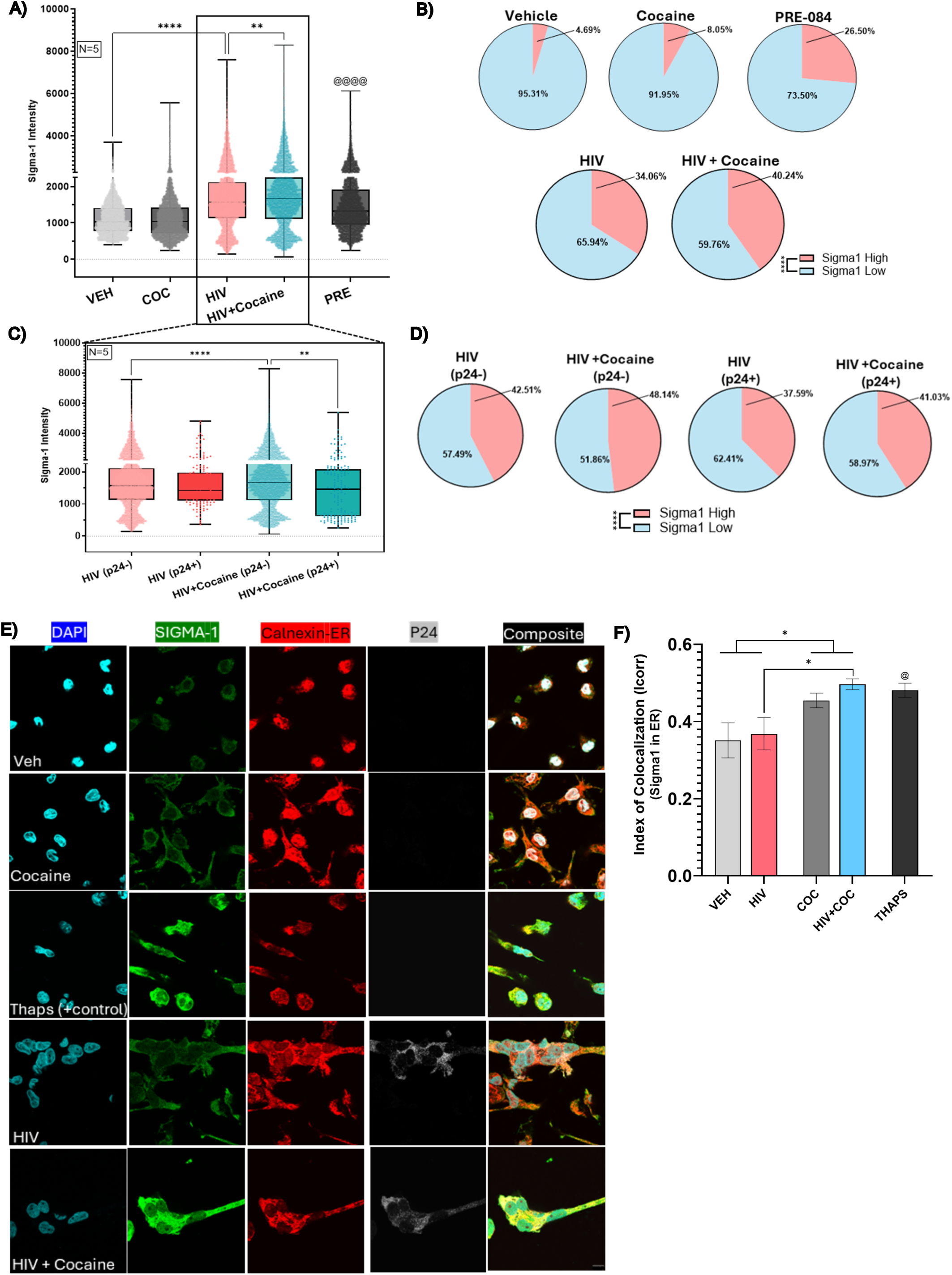
Cocaine increases the expression of sigma-1 in HIV infected cultures. **(A)** Sigma-1 intensity staining of Uninfected and Infected iMg treated with vehicle (dH20), cocaine (10^-8^M), and PRE-084. Kruskal-Wallis test with Dunn’s post hoc test. VEH vs PRE-084 (positive control): Mann-Whitney test. Scale bar = 20um. **(B)** Pie charts showing the proportion of sigma-1^high^ vs. sigma-1^low^ cells between the global treatments. Chi-square contingency test (chi-square: 3258, df: 4). **(C)** Segregated sigma-1 intensity data between p24+ (infected) vs. p24-cells (uninfected) within the same HIV vs. HIV + Cocaine cultures. Kruskal-Wallis test with Dunn’s post hoc test. **(D)** Pie chart showing the proportion of sigma-1^high^ vs. sigma-1^low^ cells between segregated p24+ vs. p24-cells. Chi-square contingency test (chi-square: 48.15, df: 3). **(E)** Subcellular localization of sigma-1 in the endoplasmic reticulum (ER) using Thapsigargin (10^-6^M) as a positive control. Nucleus (Blue), Sigma-1 (Green), ER-Calnexin (Red), p24Gag (Grey). **(F)** Quantification of colocalization between sigma-1 and calnexin. Ordinary two-way ANOVA (Coc: F (1,47 = 11.03) with Fischer’s LSD post hoc test. VEH vs Thapsigargin (positive control): Unpaired t test. *P<0.05, **<0 0.005, ***P<0.0005. Scale bar = 10um.

To examine whether these changes in sigma-1 were found in the ER, uninfected and HIV-infected iMg were treated with either Veh (H_2_O) or Coc for 3 hr at 9 DPI, using a 3 hr treatment with the ER stress inducer Thapsigargin as a positive control. Cultures were stained for ER (calnexin) to as well as sigma-1, nuclei and p24 and examined for the colocalization of sigma-1 with the ER (**Fig. 6E**). There was a main effect of cocaine on sigma-1 colocalization with the ER, but no main effect of HIV alone and no interaction between HIV and Coc (**Fig. 6F**). The amount of sigma-1-ER colocalization in the HIV+Coc cultures was significantly greater than that seen in the HIV cultures, and the lack of change in response to HIV alone was surprising give the increase seen in whole cell analyses in Fig. 6A. This suggests that while HIV increases sigma-1 expression, it does not drive the translocation of sigma-1 to the ER, and conversely, cocaine alone does not increase sigma-1 expression but does promote the movement of this receptor to the ER.

## Discussion

Substance use, specifically cocaine use, is a growing health issue, particularly within vulnerable populations such as people living with HIV. A substantial amount of research has shown that cocaine use exacerbates HIV infection *in vivo* and *in vitro.* In humanized BLT mice, cocaine administration enhanced HIV infection and circulating viral load^102,103^and similar enhancement was shown in HIV-infected huPBL-SCID mice^58^. Similar trends have been reported in other human myeloid cell models, including PBMC^104,105^, monocyte-derived macrophages (MDMs)^57,106,107^, dendritic cells^108^ ^109^ and microglia^42^. The mechanism by which cocaine acts on the viral replication process remains unclear, particularly given the absence of dopaminergic neurons, but some studies suggest the effect could be mediated via the sigma-1 receptor^42,58^. These data confirm this, showing that cocaine increases HIV infection in human iPSC derived microglia-like cells (iMg) through activation of the sigma-1 receptor. Treatment with cocaine during inoculation increased the secretion of p24 across all timepoints measured and also increased the percentage of infected (p24+) iMg at day 9 post-infection. Analysis using non-linear regression confirmed that cocaine accelerates the HIV replication in iMg. The increase in p24Gag protein expression in response to cocaine suggests that cocaine enhances transcription or translation. Additional stages of the viral lifecycle may also contribute to the increased infection, as cocaine can enhance HIV entry coreceptors in human monocyte-derived macrophages^110,111^, promote proviral DNA integration in T cells^56^, and accelerate transcription and translation in multiple models^57,60,61,102,107^. However, the involvement of transcription or translation is supported by the increased p24 expression in HIV-infected iMg treated with antiretroviral therapy (bictegravir, tenofovir alafenamide, emtricitabine), as this shows increased activity in the post-integration steps of the viral replication cycle. Cocaine primarily acts acutely via inhibition of the dopamine transporter (DAT), blocking dopamine reuptake and activating of dopamine receptors on glutamatergic and GABAergic neurons^112,113^. This process also expands the volume of tissue and number of microglia exposed to elevated dopamine^114–117^, a process that is likely enhanced in PWH, as cocaine use potentiates dopaminergic dysfunction in this population^118,119^. Cocaine exposure activates microglia^120–125^, and in PWH cocaine use worsens neurocognition^19^, disrupted response inhibition^126^ and risk processing^127^, and increased corticostriatal network deficits^128^. We and others have also shown that exposure to the concentrations of dopamine induced by cocaine use (10^-6^M to 10^-8^M) increases HIV infection and inflammation of macrophages and microglia *in vitro*^62,63,83–85^. Some studies show that the effects of cocaine on microglia are only observed *in vivo*, with *in vitro* cocaine treatment unable to recapitulate the activity^121,129–131^, a finding consistent with an indirect effect of cocaine through its actions on the dopaminergic system. While dysregulation of the dopaminergic system is clearly associated with the development of neuroHIV, cocaine mediated changes in dopaminergic neurotransmission do not explain all of the effects of Coc on HIV neuropathogenesis. Specifically, the mechanisms mediating the impact of Coc on HIV infection dynamics, and whether they involve the dopaminergic system, remain unclear, and studies have shown that Coc has other mechanisms of action beyond the modulation of dopamine. A prominent alternative pathway for Coc is the interaction with sigma-1, a multifaceted protein embedded in the ER. This interaction has emerged as a focal point in understanding Coc-induced neurochemical changes^40,41^.

These data show that cocaine mediated changes in HIV infection in iMg, do not involve the dopaminergic system, and are mediated by activation of sigma-1. Specifically, we found that the sigma-1 agonists PRE-084 and Donepezil both increased HIV replication similarly to Coc, and that the effect of Coc on HIV replication was blocked by the sigma-1 antagonist BD-1063, but not by the dopamine receptor inhibitor Flupentixol. We also found that sigma-1 expression was increased by HIV, but not by Coc treatment alone, and that Coc treatment of HIV-infected cultures was synergistic and induced a significant increase in sigma-1 expression above the HIV-infected, vehicle-treated iMg. Surprisingly, the increase in sigma-1 expression in HIV+Coc cultures was driven by changes in sigma-1 protein levels, specifically in the uninfected (p24-) iMg. This was surprising, as several studies suggest that sigma-1 activation modulates inflammatory responses^132,133^, reactive oxygen species and cell survival^49,134,135^, and we had hypothesized that sigma-1 activity also drove viral replication in the iMg. These findings contribute to a longstanding question in the field: how substance use comorbidities influence persistent neuropathology despite effective ART and suppressed viral load^11,136–138^. While some studies have attributed these residual effects to ART toxicity, our data suggest an alternate interpretation. The changes in sigma-1 within uninfected (p24-) iMg in HIV infected cultures suggest that uninfected cells exposed to HIV, although not productively infected, may play an active role in propagating or expanding chronic infection. In this case, it is possible that cocaine mediated activation of sigma-1 could have multiple effects on HIV infection, potentially increasing the susceptibility of uninfected cells to infection while also acting on post-integration processes in the infected iMg.

Sigma-1 is a multi-faceted protein involved in a large number of cellular processes, many of which could impact different stages of the viral replication cycle. As a result, scRNAseq was used to streamline pathways dysregulated by HIV + cocaine. Gene set enrichment analyses of HIV and HIV+Coc iMg found genes related to UPR, Interferon Alpha Response, DNA repair, Reactive Oxygen Species (ROS), and E2F targets as dysregulated in response to cocaine treatment. These listed pathways have been implicated in the enhancement and modulation of HIV viral pathogenesis. HIV viral proteins like Vpu modulate DNA repair mechanisms (e.g., RAD, FEN1) to suppress innate response against HIV^139,140^. Likewise, it is well established that ROS is involved in HIV pathogenesis and disease progression, with overproduction of ROS promoting HIV-associated oxidative stress^141^. Notably, the UPR, which is implicated in HIV pathogenesis and sigma-1 mediated signaling, was one of the top altered pathways, warranting further investigation.

Using RT-qPCR and Western Blotting, we found that cocaine specifically enhances signaling of the XBP1 arm of ER stress. Studies largely in T cells have found that HIV activates multiple arms of the UPR, indicating a mechanism evolved to hijack UPR signaling for its benefit^50,142,143^. Independently, cocaine enhances ER stress protein expression and sigma-1 redistribution^144–148^. Our study identified that the cocaine-mediated increase in HIV replication is at least in part driven by XBP1-mediated unfolded protein response. There was also a downregulation of host immune responses against HIV, as viral restriction factors and antiviral cytokines were reduced with cocaine treatment. Specifically, treatment reduced the secretion of IL-16, IL-12, and IL-6, similar to what is recorded in patients^37,95,96^. IL-16 and IL-12 suppress HIV replication by inhibiting viral entry^37,149,150^ and enhancing antiviral responses^151,152^ against HIV. Cocaine treatment led to increased IL-4 secretion, a cytokine known to be dysregulated during HIV infection^153^. Interestingly, IL-4 also regulates XBP1 expression, while XBP1, in turn, promotes IL-4 secretion^99^. These findings provide evidence that cocaine leverages XBP1-driven pro-viral mechanisms and immune evasion strategies to facilitate and amplify HIV infection.

These studies employed human induced pluripotent stem cell-derived microglia, which have recently emerged as a highly translatable and tractable model for investigating microglia biology^154–156^. These cells offer several advantages, including standardized and scalable generation, high phenotypic and transcriptomic similarity to primary microglia, and the presence of a human-specific immune profile not found in rodents. Further, this model is well-suited for reductionist analysis of HIV infection dynamics, inflammation, and the impact on glial health in response to HIV and/or cocaine. However, this model system also presents certain limitations. Since iPSCs are derived from a single donor, the use of a single line can limit the generalizability of findings and overlook inherent human biological variability. This is a common limitation in iPSC studies, due to the time and resources required to source and generate multiple iPSC lines. We have attempted to address this challenge by running our studies in four distinct iPSC donor lines, which is critical in the context of HIV research, where there is significant heterogeneity in infection susceptibility, disease progression, and treatment response^72,73^. While this enabled us to more effectively simulate inter-individual variability, the use of cells from only 4 individuals, as well as the variability between those lines, does present an important limitation of this work. Another caveat is that monoculture of microglia does not fully recapitulate the complexity of the brain’s microenvironment, which consists of various cell types and intricate connective circuitry that could influence the effects of HIV and/or cocaine. While we believe that studies of this type are essential to identify target genes or mechanisms that can then be evaluated in more complex co-culture systems, the results of monoculture studies should be considered carefully as they could be distinct from results found in co-cultures or *in vivo*. Finally, it is important to note that these studies did not directly assess the level of susceptibility to HIV infection in the uninfected (p24-) cells, and this is an exciting avenue for further investigation.

Collectively, these data highlight a mechanism by which cocaine enhances HIV infection of human microglia via sigma-1, perhaps in part through sigma-1 activation of the XBP-1 mediated UPR pathway. Our scRNAseq data showed a number of potential other pathways, including DNA repair, oxidative phosphorylation, reactive oxygen species pathway, etc. by which cocaine mediated sigma-1 activation could also influence HIV replication and persistence in the CNS. Notably, these effects may be mediated not only by productively infected cells but also by bystander cells exposed to HIV and cocaine. Future investigations into how bystander microglia populations contribute to neuropathogenesis may uncover new understandings of HIV persistence. Additionally, these data suggest that targeting sigma-1 alongside ART may offer a novel therapeutic strategy for addressing comorbid cocaine use and HIV infection, which is currently under investigation^157^. These observations underscore the importance of evaluating context of the “protective” effects that have been attributed to sigma-1, as targeting this receptor may be therapeutic in some cases but damaging in others, potentially promoting viral persistence in PLWH. The data also raises important considerations for other conditions and treatments that engage this pathway. Donepezil and other Alzheimer’s disease treatments, antidepressants such as fluvoxamine (Luxor) and sertraline (Zoloft), and a number of other therapeutics either on the market or in clinical trials, all shown agonism at sigma-1^158–160^. These findings identify sigma-1 as a key link between HIV-associated cellular stress and cocaine exposure, highlighting shared mechanisms that could inform future therapies not only for HIV neuropathogenesis but also for broader treatment of SUD and neuropsychiatric disorders.

## Acknowledgements

We would like to thank all members of the Gaskill, Matt, and Fox Labs who contributed to this work with their feedback and discussion. We would also like to thank the Human Pluripotent Stem Cell Core, specifically Dr. Deborah French and Dr. Jean-Anne Maguire, for providing the common myeloid progenitors and assisting with the development of our iMicroglia model.

## Funding

This work was supported by grants from the National Institutes of Drug Abuse, DA057337 and DA058051 (PJG), MH132466 (SMM), U01 DA053624 (HF) and the Department of Pharmacology and Physiology at Drexel University College of Medicine.

## Competing interests

The authors report no competing interests.

## Supplementary material

Supplementary material is available at *Brain* online.

## Notes

### Competing Interest Statement

The authors have declared no competing interest.

