## Supplementary Methods for "Cocaine Acts Through Sigma-1 to Enhance HIV Infection of Microglia"

### Supplemental Methods

#### Reagents

RPMI-1640 (cat # 11875119) medium, penicillin/streptomycin (P/S) (cat # 15140163), TrypLE Express (cat # 12604013) were from Invitrogen (ThermoFisher, Carlsbad, CA, USA). Bovine serum albumin (BSA) (cat # BP1600100), glycine (cat # G48500), and 4% Paraformaldehyde solution (cat # AAJ19943K2) were from Fisher Scientific (Waltham, MA, USA). Fetal calf serum (FBS) was from Corning (cat # MT35010CV). Hydroxyethyl piperazineethanesulfonic acid (HEPES) (cat # BP2991), Tween (cat # NC0689073), and dimethyl sulfoxide (DMSO) (cat # 472301) were obtained from Sigma-Aldrich (St. Louis, MO, USA). Macrophage colony-stimulating factor (M-CSF) (cat # 300-25), IL-34 (cat # 200-34), and TGF-β1 (cat # 100-21) were from Peprotech (Rocky Hill, NJ, USA). Cocaine hydrochloride (DEA Code 9041) was from the NIDA Drug Supply Program (NDSP). PRE-084 (cat # 9001329), Bictegravir (cat # 26532), Tenofovir Alafenamide (cat # 26214), and Emtricitabine (cat # 16879) were from Cayman Chemical Company (Ann Arbor, MI, USA). Donepezil (cat # 4385) and BD-1063 (cat # 0883) are from Tocris Bioscience (Bristol, UK). Flupentixol dihydrochloride (cat # 4057/50) was from R&D Systems (Minneapolis, MN, USA). All drugs were resuspended in their respective solvents, with Biktarvy (bictegravir, emtricitabine, and tenofovir alafenamide) in DMSO and others in dH2O, then aliquoted and stored in -20°C until use.

TaqMan Fast Universal Master Mix, and PCR assay probes for SIGMAR1 (Hs00195337_m1), XBP1 (Hs00231936_m1), ATF6 (Hs00231758_m1), ATF4 (Hs00909568_g1), CHOP (Hs00358796_g1), IFITM1 (Hs00705137_s1), IFIT1 (Hs01675197_m1), IFIT2 (Hs00533665_m1), MALAT1 (Hs00273907_s1), and STMN1 (Hs01027515_gH) and 18s (4319413E) genes were purchased from Applied Biosystems (ThermoFisher, Waltham, MA, USA).

#### Generation of $HIV_{ADA}$ Viral Stocks

Viral stocks were generated by infecting CEM-SS cells with a blood-derived, R5-tropic strain of HIV ($HIV_{ADA}$[1]), as we have done previously [2,3]. Briefly, CEM-SS cells were cultured in

T175 flasks in RPMI-1640 containing 10% FBS, 1% PenStrep, and HEPES. Each culture was inoculated with a single tube of $HIV_{ADA}$ from a previous stock, with media changes every 3 – 4 days. Initial $HIV_{ADA}$ stocks were obtained through the NIH HIV Reagent Program, Division of AIDS, NIAID, NIH: Human Immunodeficiency Virus-1 ADA, ARP-416, contributed by Dr. Howard Gendelman. Infected cultures were maintained until syncytia formation was observed, generally after 15 – 18 days. Once syncytia were observed, supernatant was collected every 24 hours until viral replication outpaced cell growth, killing the culture. The stocks used in the study showed syncytia at 17 days post-infection, and cell-free supernatants were collected daily from 18 to 41 days post-infection. To collect supernatants, cultures were centrifuged at 1200 x g to pellet cells and debris, then the cell pellet was resuspended in media, transferred to a fresh conical vial, aliquoted in 1 mL aliquots, and stored at -80°C for use as viral stocks. The concentration of viral stock was determined by quantifying the amount of p24•Gag per mL, using an HIV p24 high-sensitivity AlphaLISA Detection kit (Revvity, Waltham, MA).

#### Lactate Dehydrogenase (LDH) Assay

To evaluate cell viability, iMg supernatants were quantified for LDH release using the CyQUANT$^{tm}$ LDH Cytoxicity Assay (ThermoFisher Scientific # C20301). In brief, 50ul/12.5ul of sample were loaded in 96-well/384-well plates and incubated with equal amounts of reaction mixture (LDH substrate mix + assay buffer) for 30 minutes at 25°C in the dark. An equal amount stop solution was added to each well after the incubation and absorbance values at 490nm (signal) and 680nm (background) were taken on a spectrophotometer. A positive control included in the kit to verify assay was successful and media control was used to quantify spontaneous LDH release. Absorbance (viability) was calculated by calculating the using the equation: Sample absorbance - Spontaneous LDH release.

#### Quantitative Polymerase Chain Reaction

Infected mature iMg cultured in 24-well plates (P24-1.5P, Cellvis) were treated with vehicle ($H_2O$) or cocaine ($10^{-8}$M) for 3 hours. Total RNA was collected using the RLT buffer (74134, Qiagen), and RNA extraction was performed using the RNeasy Plus kit (74134, Qiagen) according to the manufacturer's protocol. RNA quantity and purity were determined by a NanoDropOne spectrophotometer (Nanodrop Technologies). RNA (200-250ng) was used to synthesize cDNA using the high-capacity reverse transcriptase cDNA synthesis kit (4368814, Abcam). Quantitative PCR (qPCR) was performed on a QuantStudio 7 (Thermo Fischer) using gene-specific, Taqman primers obtained from Thermo Fischer. The genes SIGMAR1, XBP1, ATF6, ATF4, CHOP, IFITM1, IFIT1, IFIT2, MALAT1, and STMN1 were amplified from cDNA by quantitative PCR. The gene 18s was used as a housekeeping gene during all qPCR assays.

#### Multiplex Cytokine Analysis

For cytokine analysis, supernatants were assessed using the MSD V-Plex Human Neuroinflammation panel (Cat #K15210D-1, Meso Scale Diagnostics, LLC, Rockville, Maryland, USA). Biomarkers analyzed using this assay included IL-16, IL-12, IL-6, IL-7, and IL-4. Samples were run in duplicate according to the manufacturer's protocol. All plates were imaged on the MESO QuickPlex SQ 120MM and analyzed using MSD Discovery Workbench 4.0 Software (Meso Scale Diagnostics, LLC, Rockville, Maryland, USA).

#### Immunofluorescence

iMg plated in 96 wells at day 9 post-infection were fixed with 4% PFA at 4°C for 20 minutes, permeabilized, and incubated with blocking buffer (1% BSA, 0.1% Tween 20, and 22.52 mg/mL glycine in 1X PBS) for 1 hour at room temperature. For 3-color immunofluorescence staining of p24+ cells, cells were incubated with anti-HIV-1 p24 monoclonal (AG3.0) primary antibody (4 µl/ml, NIH HIV Reagent Program catalog # 4121) overnight at 4°C. After the primary incubation, the cells were incubated with Alexa Fluor 488 secondary antibody (A-11001, Thermo Fisher) for 1 hour at room temperature. The cells were counterstained with DAPI (0.2 µg/mL, D1306, Thermo Fisher) and CellMask Deep Red, CMDR (250ng/mL, C10046, Thermo Fisher) for 10 minutes, then preserved in 1x PBS.

For 4-color immunofluorescence staining of sigma-1, infected iMg cells were treated with vehicle (H20), cocaine ($10^{-8}$M), PRE-084, or Thapsigargin (positive control, $10^{-6}$M) for 3 hours to examine changes in sigma-1 before fixation and blocking. Supernatants were collected for cytokine analysis. Cells were incubated with anti-HIV-1 p24 monoclonal (AG3.0) primary antibody (4 µl/ml, NIH HIV Reagent Program catalog # 4121) and anti-SIGMAR1 polyclonal antibody (1:50, 15168-1-AP, Proteintech) overnight at 4°C. After the primary incubation, the cells were incubated with Alexa Fluor 488 secondary antibody (A-11008, Thermo Fisher) and DyLight

755 secondary antibody (SA5-10175, Thermo Fisher) or Alexa Fluor 555 secondary antibody (A-21422, Thermo Fisher) and Calnexin Monoclonal Antibody (AF18), Alexa Fluor 647 (MA3-027-A647, Thermo Fischer) for 1 hour at room temperature. The cells were counterstained with DAPI (0.2 µg/mL, D1306, Thermo Fisher) and CellMask Deep Red, CMDR (250ng/mL, catalog # C10046 Thermo Fisher) for 10 minutes, then preserved in 1x PBS. CMDR staining was excluded during subcellular localization experiments with Calnexin.

#### High Content Imaging

High-content imaging of the cells was performed using the Cellomics CellInsight CX7 HCS Platform, capturing 20 fields per well at 20x magnification. To analyze the percentage of p24+ cells, a 2-channel colocalization was performed using the sample parameters included in **Supplementary Table 1.** The CMDR stain (Channel 1) was used to identify cells in the well, and the anti-P24 stain (Channel 2) was used to identify p24+ cells. Both channels were gated by excluding border objects, object area, and object total/variant intensity (as needed). A region of interest (ROI) was formed based on the colocalization between CMDR and anti-p24 (Target1). Cell-level data was exported on ROI_A_ Target1_ObjectVarInten and used to calculate the %p24+ cells from the p24+ cell counts and total cell counts. Other cell morphology parameters were also exported from Channel 1 (CMDR), like perimeter-to-area (P2A) ratio and cell area.

To quantify multinucleated giant cells (MNGC; nuclei >5), a 2-channel colocalization was performed using the specific parameters included in **Supplementary Table 2.** The CMDR stain (Channel 1) was used to identify cells, and the DAPI stain (Channel 3) was used to identify nuclei. Both channels were gated by object area and object total/variant intensity (as needed). A region of interest (ROI) was formed based on the colocalization between CMDR and DAPI (Target1). Cell-level data was exported on ROI_B_Target_I_ObjectCount and used to calculate the % MNGC cells from which an Infected MNGC index was calculated (% MNGC/%Infected Cells).

For sigma-1 imaging, the images were analyzed using the HCS Studio Software, specifically a 3-channel colocalization using the sample parameters included in **Supplementary Table 3.** Each channel was gated by excluding border objects, object area, and object total/variant intensity (as needed). Two regions of interest were formed based on the colocalization between CMDR and anti-p24 (ROI A) and CMDR and anti-SIGMAR1 (ROI B). Cell-level data was exported on each ROI and filtered to identify overall sigma-1 intensity between conditions and the sigma-1 intensity between the p24+ vs p24- cells. For subcellular localization studies, two to four 60X images per well were acquired using an Olympus FV3000 confocal laser scanning microscope. Field-level images were analyzed using the ImageJ plugin "Colocalization ColorMap" to quantify colocalization between ER (Calnexin) and Sigma-1. High-resolution representative images were acquired at 60X (2x zoom) using an Olympus FV3000 confocal laser scanning microscope.

#### Western Blotting

Protein samples were collected using the Qiagen AllPrep DNA/RNA/Protein mini kit (Qiagen #80004). For protein isolation, samples were incubated with ice-cold acetone (1:4 dilution) for 30 minutes, followed by centrifugation at 13,000 rpm for 10 minutes at 4°C. The resulting protein pellets were washed with molecular grade ethanol and resuspended in M-PER lysis buffer (Thermo Fisher #78501). Following sonication, protein concentrations were determined using the BCA protein assay kit (Thermo Fisher #23227). Western blotting was performed as previously described[4]. Twenty micrograms of protein were loaded into each lane of a 4-12% bis-tris plus wedge well gel (Thermo Fisher #NW04120BOX) for electrophoretic separation. Proteins were subsequently transferred to a PVDF membrane (Thermo Fisher #88518). Total protein loading was verified using Revert™ 700 Total Protein Stain (LI-COR #926-11011), which was further used for expression normalization. Following the removal of the total protein stain, membranes were blocked for 1 hour at room temperature using 5% BSA solution. Primary antibodies [Cell Signaling XBP1-s #D2C1F] were applied at a 1:1000 dilution and incubated overnight at 4°C. After three washes with TBS-T, membranes were incubated with anti-rabbit IgG HRP-linked secondary antibody (Cell Signaling #7074P2) for 1 hour at room temperature. Following three additional TBS-T washes, protein bands were visualized using a 3:1 mixture of Pico PLUS Chemiluminescent and Femto Maximum Sensitivity substrates (Thermo Fisher #34580 and #34096, respectively). Images were captured using the LI-COR Odyssey FC dual-mode western blot imaging system and analyzed with ImageStudio Lite software.

#### Single-cell RNA sequencing

For alignment, the HIV genome was modified to exclude one of the long terminal repeats (3' LTR). Summary statistics from the Cell Ranger gene counting output are provided in **Supplementary Table 4**. All downstream analyses were performed in Python (v3.12.7) using Scanpy (v1.10.3)[5]. Raw count matrices from vehicle- and cocaine-treated samples were imported and merged. Features expressed in at least three cells were retained. Initial quality control (QC) involved filtering cells based on unique molecular identifiers (UMIs), gene counts, and mitochondrial transcript percentage. Following QC, data were normalized using sc.pp.normalize_total and log-transformed with sc.pp.log1p. Highly variable genes were identified using sc.pp.highly_variable_genes and subsequently scaled. To account for technical variation, total counts per cell and mitochondrial percentage were regressed out using sc.pp.regress_out. Principal component analysis (PCA) was performed via sc.tl.pca, and the first 15 principal components (PCs) were used to construct the neighborhood graph with sc.pp.neighbors. Clustering was performed using the Leiden algorithm (sc.tl.leiden) with the flavor set to "igraph", n_iterations = 2, and resolution = 0.5. Uniform manifold approximation and projection (UMAP) was used for dimensionality reduction and visualization (sc.tl.umap). Cluster-specific differentially expressed genes (DEGs) were identified using the Wilcoxon rank-sum test implemented in sc.tl.rank_genes_groups. DEGs were filtered by adjusted p-value < 0.05 and absolute log2fold change > 0.5.
