## Supplementary Figures for "Cocaine Acts Through Sigma-1 to Enhance HIV Infection of Microglia"

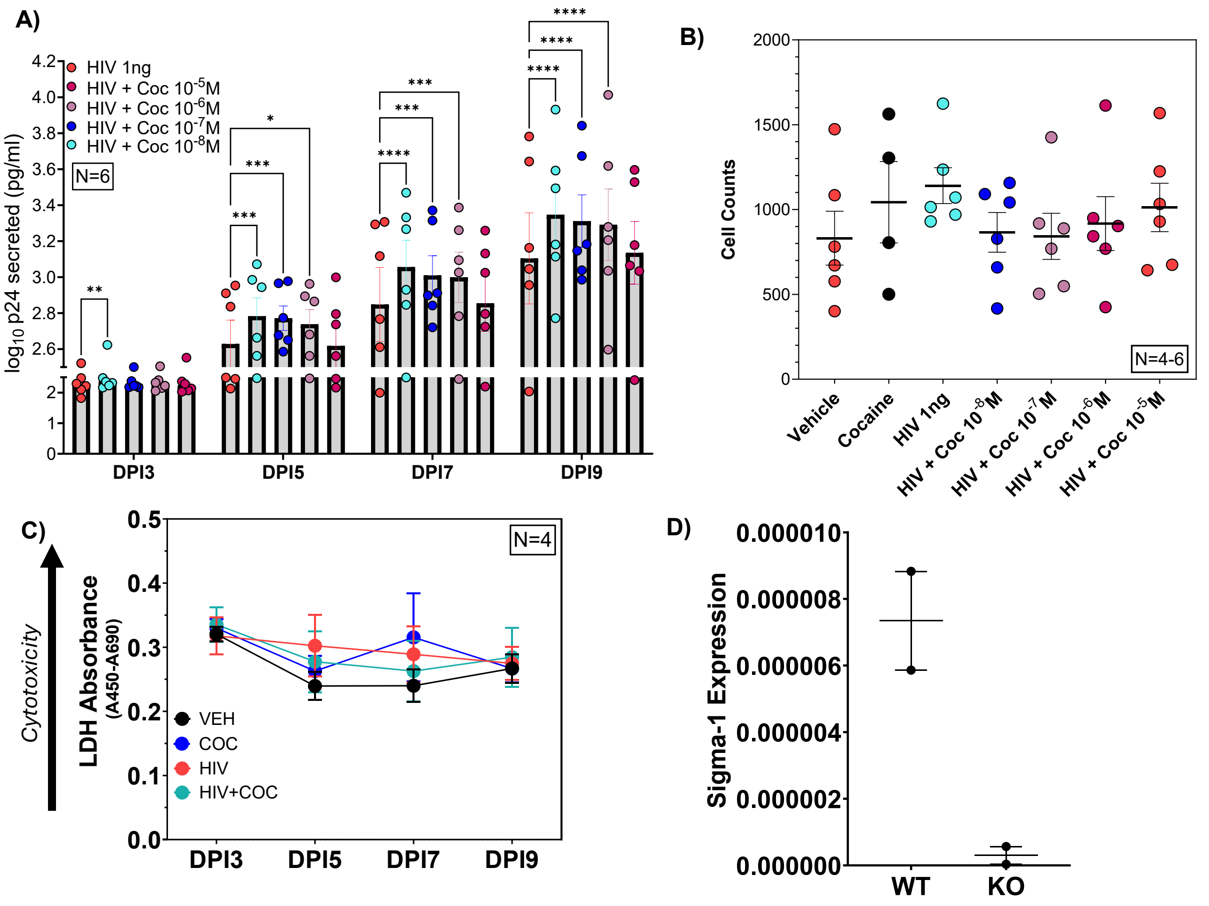


**Supplementary Figure 1: Validating Cocaine HIV Infection Curve and Cytotoxicity. (A)** p24AlphaLISA of iMg treated with HIV +/- cocaine ($10^{-5}$M – $10^{-8}$M) from DPI3 -DPI9. N=6. Two-way ANOVA ($F[3,15] = 40.52$, Dunnett's post hoc test. ****$P<0.0001$, ***$P<0.001$, **$P<0.01$, *$P<0.05$. **(B)** iMg cell count at DPI9. **(C)** LDH Cytotoxicity assay measuring cell death following HIV+/-Cocaine ($10^{-8}$M).

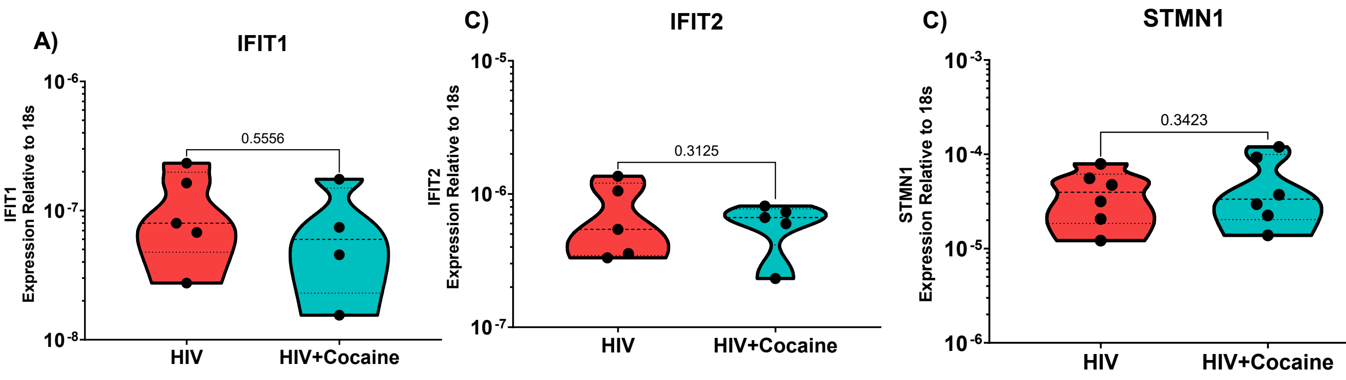


**<u>Supplementary Figure 2:</u> Effect of cocaine on innate immune and unfolded protein response.** qPCR Gene expression validation of more **(A-B)** innate immune response genes and **(C)** HIV transcriptional regulation gene. Paired and unpaired t-test.

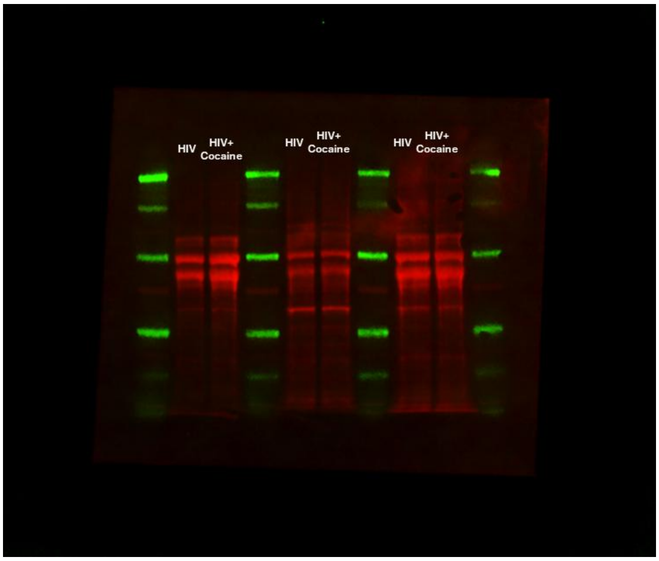


Total Protein Stain

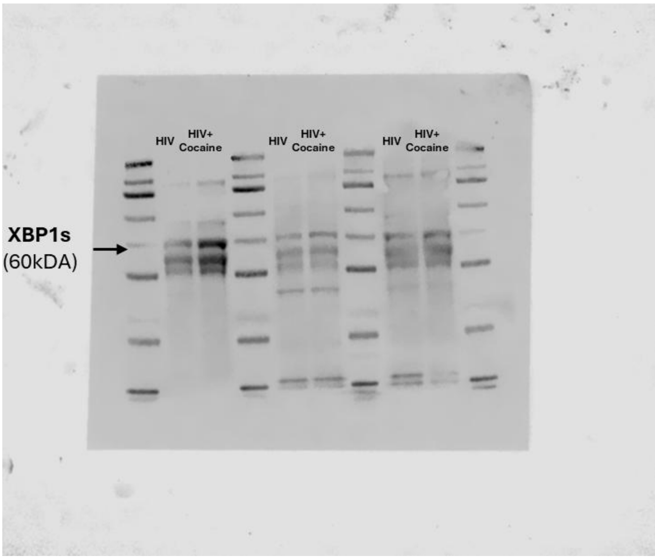


XBP1s

**<u>Supplementary Figure 3:</u> Full Western Blots for XBP1s and Total Protein Stain**
